# Hamstrings are stretched more and faster during accelerative running compared to speed-matched constant speed running

**DOI:** 10.1101/2024.03.25.586659

**Authors:** Reed D. Gurchiek, Zachary Teplin, Antoine Falisse, Jennifer L. Hicks, Scott L. Delp

**Affiliations:** Department of Bioengineering, Clemson University, Clemson, SC 29634, USA; Department of Bioengineering, Stanford University, Stanford, CA 94305, USA; Department of Mechanical Engineering, Stanford University, Stanford, CA 94305, USA

## Abstract

**Background:** Hamstring strain injuries are associated with significant time away from sport and high reinjury rates. Recent evidence suggests that hamstring injuries often occur during accelerative running, but investigations of hamstring mechanics have primarily examined constant speed running on a treadmill. To help fill this gap in knowledge, this study compares hamstring lengths and lengthening velocities between accelerative running and constant speed overground running.

**Methods:** We recorded 2 synchronized videos of 10 participants (5 female, 5 male) during 6 accelerative running trials and 6 constant speed running trials. We used OpenCap (a markerless motion capture system) to estimate body segment kinematics for each trial and a 3-dimensional musculoskeletal model to compute peak length and step-average lengthening velocity of the biceps femoris (long head) muscle-tendon unit. To compare running conditions, we used linear mixed regression models with running speed (normalized by the subject-specific maximum) as the independent variable.

**Results:** At running speeds below 75% of top speed accelerative running resulted in greater peak lengths than constant speed running. For example, the peak hamstring muscle-tendon length when a person accelerated from running at only 50% of top speed was equivalent to running at a constant 88% of top speed. Lengthening velocities were greater during accelerative running at all running speeds. Differences in hip flexion kinematics primarily drove the greater peak muscle-tendon lengths and lengthening velocities observed in accelerative running.

**Conclusion:** Hamstrings are subjected to longer muscle-tendon lengths and faster lengthening velocities in accelerative running compared to constant speed running. This provides a biomechanical explanation for the observation that hamstring strain injuries often occur during acceleration. Our results suggest coaches who monitor exposure to high-risk circumstances (long lengths, fast lengthening velocities) should consider the accelerative nature of running in addition to running speed.

## 1. Introduction

Hamstring strain injuries are a major time-loss injury for athletes in many sports including soccer^1^, track and field^2^, basketball^3^, American football^4^, and rugby^5^. A recent meta-analysis^6^ of 63 articles and more than 7 million exposure hours estimated that hamstring strains comprised 10% of all injuries in field-based sports. This analysis revealed relatively constant injury rates over the last 30 years, while Ekstrand et al. reported increases in elite soccer^7^. Reinjury rates are high^8,9^. Injury prevention is thus paramount, but knowledge about injury mechanisms is still limited. A better understanding of the movement conditions that pose a high risk for injury could lead to innovations in preventative screening and athlete monitoring, and interventions to prevent injury.

The peak length of the hamstring muscle increases with running speed both at the muscle fiber^10^ and muscle-tendon unit^11^ levels. High running speeds also present large hamstring lengthening velocities under high loads^12^. These eccentric loads are associated with increased muscle fiber strains^13^ and eccentric work, predictors of microdamage^14–16^. These biomechanical features of the hamstrings during high-speed running (large peak lengths, lengthening velocities, and eccentric loads) are thought to contribute to injuries^12,17,18^ and may help explain the occurrence of injuries during high-speed running^19–22^. As a result, high-speed running is often used as an index of load for athlete monitoring^23,24^.

The eccentric loading and injury incidence associated with running has motivated the investigation of hamstring mechanics during high-speed running^25^. For example, kinematic analyses have shown that hamstring lengths (normalized by the length in an upright posture) peak in late swing phase and are greatest in the biceps femoris long head compared to the other hamstring muscles due to moment arm differences^11,26^. Kinetic analyses have demonstrated that hamstring loads may exceed 2000 N^27^ and are greater in swing phase (10.5-26.4 N/kg) than in stance phase (4.6-11.6 N/kg)^25^. Moreover, eccentric work is done exclusively in swing phase^26,28^ and hamstring muscle activity during stance phase is greatest at foot contact^29^.

Previous investigations of hamstring injury mechanisms have primarily considered running at relatively constant and high speeds^25,30^. However, it was recently shown that hamstring injuries often occur when athletes are accelerating^31^ and not necessarily at high running speeds^22^.

Moreover, the increase in injury incidence in elite soccer players reported by Ekstrand et al. between 2001-2014^32^ may partly result from an evolution in the nature of running activity soccer players perform. For example, the number of sprints, sprint distance, and “explosive” (high-acceleration) sprints performed by soccer players all increased within a similar time frame (2006-2013)^33^.

While the biomechanics of accelerative running have been explored^34–37^, the kinematics of the hamstrings (muscle-tendon length and lengthening velocity) during acceleration are less understood. The hamstrings are important for generating propulsive ground reaction forces during acceleration^35,36^, with large hamstring muscle forces (6-8 bodyweights) during a sprint that starts from stationary^35^. In a prospective study, Schuermans et al. (2017) identified increased anterior pelvic tilt and increased frontal plane trunk bending as kinematic features during the acceleration phase of sprinting of soccer athletes who sustained an injury compared to those who did not^38^. Higashihara et al. (2018) found the biceps femoris was more active than the semitendinosus during stance phase in accelerative running, whereas the semitendinosus was more active in swing phase in both accelerative and top speed running^39^. However, this study considered accelerative running at 15 m after the start of the sprint wherein athletes had already obtained a relatively high speed (an average of 8.54 m/s, approximately 90% of the average top speed of 9.52 m/s). Schuermans et al. (2017) observed a similar phase of the sprint (15-25 m after the start), but running speeds were not reported. Thus, the effect of running speed on hamstring mechanics (i.e., both kinematics and kinetics) during accelerative running remains unclear. Furthermore, the mechanics of a runner accelerating from a relatively high speed may not closely represent accelerative running activity during gameplay, which often is initiated from submaximal running speeds (a “flying” sprint). Finally, hamstring lengths and lengthening velocities during accelerative running have not been explored using a 3-dimensional musculoskeletal model similar to the work of Thelen et al. (2005) for non-accelerative, high-speed running on a treadmill^11^.

Given the lack of knowledge about hamstring lengths and lengthening velocities during accelerative running, we aimed to compare these variables during accelerative running to running at a constant speed across a range of speeds. To emulate in-game running activity, we used a markerless motion capture technology (OpenCap^40^) that allowed analysis of overground accelerative running in an unconstrained outdoor setting. We hypothesized the hamstrings would be subject to greater peak muscle-tendon lengths and lengthening velocities in accelerative running compared to constant speed running.

## 2. Material and Methods

### 2.1 Data Collection

Ten participants volunteered to participate in this study (Table 1). They were recreationally active with an average self-reported exercise frequency of 5 days per week for 1 hour per day. Following a self-directed warm-up, participants performed a 30-meter maximal effort sprint, 6 constant speed running trials, each at a different speed, and 6 accelerative running trials. The 30-meter maximal effort sprint was used to determine the subject-specific top running speed. The accelerative running trials were flying sprints wherein participants began running at a constant speed and, when given a verbal cue, accelerated forward as fast as possible. These trials were designed to represent in-game accelerative running activity in team sports where sprinting is often initiated from a moving start. The constant speeds that preceded the 6 accelerations were the same as those in the constant speed running trials. The first two running speeds were self-selected (jogging and brisk running) and the last 4 were prescribed (3, 4, 5, and 6 m/s, performed in that order). At each speed, subjects performed the constant speed running trial first followed by the accelerative running trial and were given as much rest time between trials as desired to avoid fatigue. The order of running speeds and conditions was not randomized. All trials were performed overground on an outdoor running surface. All study activities were approved by the Stanford University Institutional Review Board. Informed written consent was obtained from all study participants.

**Table 1.**
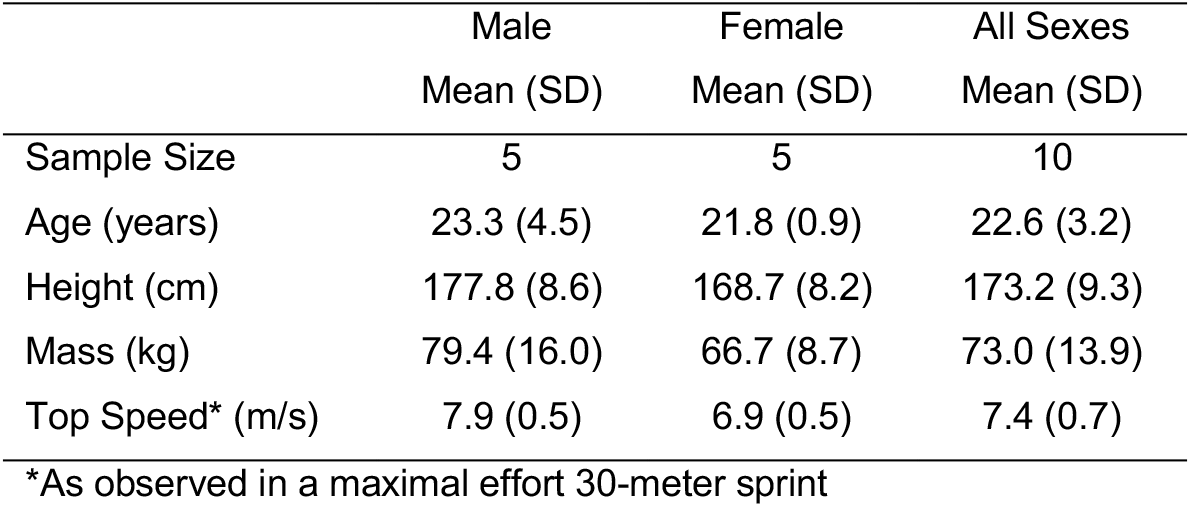
Subject characteristics.

|  | Male | Female | All Sexes |
| --- | --- | --- | --- |
|  | Mean (SD) | Mean (SD) | Mean (SD) |
| Sample Size | 5 | 5 | 10 |
| Age (years) | 23.3 (4.5) | 21.8 (0.9) | 22.6 (3.2) |
| Height (cm) | 177.8 (8.6) | 168.7 (8.2) | 173.2 (9.3) |
| Mass (kg) | 79.4 (16.0) | 66.7 (8.7) | 73.0 (13.9) |
| Top Speed* (m/s) | 7.9 (0.5) | 6.9 (0.5) | 7.4 (0.7) |

\*As observed in a maximal effort 30-meter sprint

For the jogging self-selected speed, participants were instructed to run at a comfortable jogging speed. For the brisk-running self-selected speed, they were instructed to run at a brisk running speed that was faster than the jogging speed, but not a maximal effort sprint. Participants matched prescribed speeds by self-modulating their running speed such that they passed cones positioned along the running lane at uniform distances (2.5 or 5.0 meters) to the beat of a metronome. The metronome frequency was based on the distance between cones and the prescribed speed (e.g., 60 beats per second with 5-meter cone spacing for a 5 m/s speed prescription). Subjects practiced running at each speed before data was collected. It was not imperative that speeds were matched exactly, but that a range of speeds were captured to allow investigation of the effects of running speed.

Two mobile phones were used to record videos (60 frames per second) of the running trials. The phones were attached to tripods placed such that the center of the capture volume was approximately 35 meters from the start of all running trials. Each phone was positioned 4 meters from this 35-meter mark at an angle from the direction of travel on each side of the running lane ensuring both cameras could view the anterior-frontal plane along with the right-sagittal plane for one and the left-sagittal plane for the other.

### 2.2 Data Analysis

Video recording, synchronization, and processing as well as the scaling and kinematic analysis of a 3-dimensional musculoskeletal model^41^ were performed with OpenCap^40^ using OpenPose^42^ as the underlying pose estimation model. We reprocessed the video data to leverage OpenPose’s high accuracy settings, which were shown to provide more accurate results than the default settings at the expense of computational cost^40^. The model was scaled using video data from a static standing calibration trial. We identified consecutive foot contact events to segment kinematic data into individual steps. Foot contact events were identified as the instant coincident with the negative-going zero crossing in the shank sagittal plane angular velocity (lowpass filtered with a 2 Hz cutoff frequency for event detection only). This event follows a relatively large-magnitude, positive signal associated with the swing phase^43^ and thus is reliably identified algorithmically and facilitates a consistent step segmentation. We analyzed the 2 steps (1 full gait cycle) immediately following the initiation of acceleration in the accelerative running trials and the 3 steps closest to the center of the capture volume in the constant speed trials.

The length of the biceps femoris (long head) muscle-tendon unit (MTU) was calculated using OpenSim^44^ and the subject-specific musculoskeletal model. The MTU length is dependent only on the hip and knee joint angles, which were calculated within OpenCap and were lowpass filtered with a zero-phase shift, 4th order Butterworth filter with 15 Hz cutoff frequency adjusted for the backward pass^45^. The 15 Hz cutoff frequency was chosen because, on average, 90% of the hip and knee flexion angle signal power was contained below 15 Hz across all running trials. Hip and knee flexion speed were computed by numerically differentiating the hip and knee flexion angles with respect to time using the 5-point central difference method. The MTU lengthening velocity was calculated in OpenSim using the joint angles and speeds. For each step from each trial, we used the peak MTU length and average lengthening velocity (on the side exclusively in the swing phase during the step) as primary outcomes to compare running conditions. We chose to explore the average lengthening velocity instead of the peak because it is more robust to noise introduced from numerical differentiation. Nonetheless, we also explored the peak lengthening velocity in a post-hoc analysis (results reported in the supplementary material).

### 2.3 Data Quality Assessment

We identified and removed 61 poorly reconstructed steps (out of 300 total) leaving 239 available for analysis (140 steps for constant speed running and 99 steps for accelerative running). This was based on several data quality checks. First, trials were removed if the kinematic reconstructions were unacceptable following visual inspection in the OpenSim graphical user interface or if any foot contacts were not identified. Then, an individual step was removed if either outcome measure (peak length or lengthening velocity) for the step was deemed an outlier, called a discrete outlier, or if one of several other relevant kinematic time-series for the step was deemed an outlier, called a time-series outlier. A discrete outlier was defined as a value that was greater than 3 standard deviations away from the mean. For detection of time-series outliers, all time-series were first resampled to express their values as a function of the percentage of the step duration. A time-series was labeled an outlier if any of these discrete-time values met the discrete outlier criterion. The time-series included for outlier detection were the hip and knee flexion angle and speed and the MTU length and lengthening velocity. The mean and standard deviation used for outlier detection were computed using data from all participants, but specific to running condition (constant speed or accelerative) and running speed (0-55%, 55-65%, 65-75%, 75-85%, and 85-100% of top speed).

### 2.4 Statistical Analysis

We used 2 separate linear mixed model regression analyses to compare the primary outcomes (MTU peak lengths and MTU lengthening velocities) between running conditions with running speed as the independent variable. To account for size variations across participants, we normalized MTU lengths and lengthening velocities by the MTU length in the neutral configuration (hereon referred to as the neutral length). To account for variability in top running speed across participants, we normalized running speeds by the subject-specific maximum observed in the 30-meter sprint. Each regression model had 4 fixed-effects coefficients including 2 condition-specific intercepts and 2 condition-specific slopes (modeling the effect of running speed) along with 4 random effects (condition-specific slopes and intercepts) per participant. Running speeds were shifted (normalized running speed minus 1) for model fitting such that the intercept corresponded to the response (peak length or lengthening velocity) at top speed. We used the fitlme function in Matlab (R2021a, The MathWorks, Inc., Natick, MA) for model fitting using restricted maximum likelihood estimation (REML) and estimated all elements of the covariance matrix (i.e., a FullCholesky covariance pattern).

To explore the effects of different joint and segment kinematics on the primary outcomes, we also fit 6 additional linear mixed regression models with the same structure as for the primary outcomes, but with different response variables: hip flexion, knee flexion, pelvic tilt, and thigh tilt (relative to the global vertical axis) angles at the instant of peak MTU length and the contributions of the hip and knee flexion speed to the MTU lengthening velocity. The latter were determined by first computing the product of the hip or knee flexion speed and the joint angle specific MTU moment arm. The sample average across all products for which the MTU was lengthening was used as the response. Pelvic and thigh tilt angles were calculated using an Euler angle decomposition (ZXY for the pelvis, ZYX for the thigh) of the orientation (rotation matrix) of each segment relative to the global coordinate system. The tilt angle for both segments was the angle associated with the first rotation (about the Z-axis). Because the motion is primarily sagittal, the tilt angle can be interpreted as the angular deviation of the segment long axis relative to the global vertical axis. For example, the pelvic and thigh tilt angle would both be 0° during a standing posture, 90° during a supine posture, and -90° during a prone posture.

## 3. Results

The general pattern of the biceps femoris MTU length throughout the step was similar for constant speed and accelerative running (Figure 1). Minimum MTU length occurred at approximately 25% of the step cycle. The hamstrings then lengthened up to a peak length at approximately 75-80% of the step cycle and terminated with a brief shortening phase just before foot contact (Figure 1, top row).

**Figure 1.**
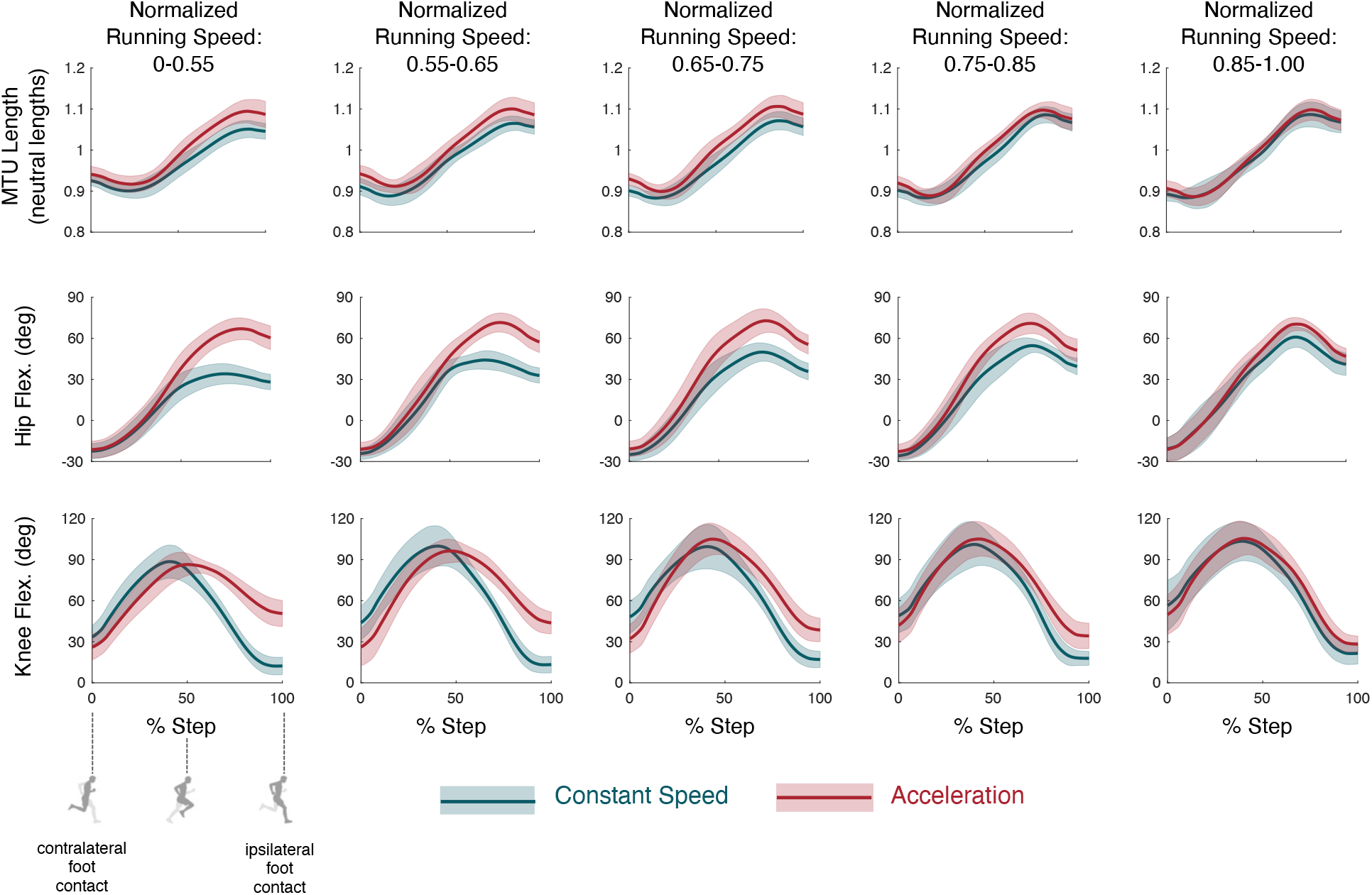
Biceps femoris MTU length (top row, units: neutral lengths), hip flexion angle (middle row, units: degrees), and knee flexion angle (bottom row, units: degrees) as a percentage of the step duration (contralateral foot contact to ipsilateral foot contact) for the accelerative (red) and constant speed (teal) running conditions. Solid lines and shaded areas indicate the ensemble mean +/- standard deviation across all participants within the same speed bin (indicated at the top of each column). Running speeds are normalized by the subject-specific top speed.

Despite similarity in the general pattern of the MTU length trajectories, accelerative running resulted in greater peak MTU lengths, with more pronounced differences for slower running speeds (Figure 1). At top speed, the peak lengths were 111-112% of neutral length (11-12% strain), and this was not different between conditions (p=0.431, Figure 2A). However, the peak MTU length was greater (p<0.001, Figure 2A) when accelerating from slow speeds than in speed-matched, constant speed running (i.e., below 75% of top speed based on 90% prediction intervals in Figure 2A). For example, the regression analysis predicted that the peak MTU length experienced when a person accelerates while running at only 50% of top speed is the same as if running with a constant speed of 88% of top speed. (The linear mixed regression results are also tabulated in Table S1 of the supplementary material.)

**Figure 2.**
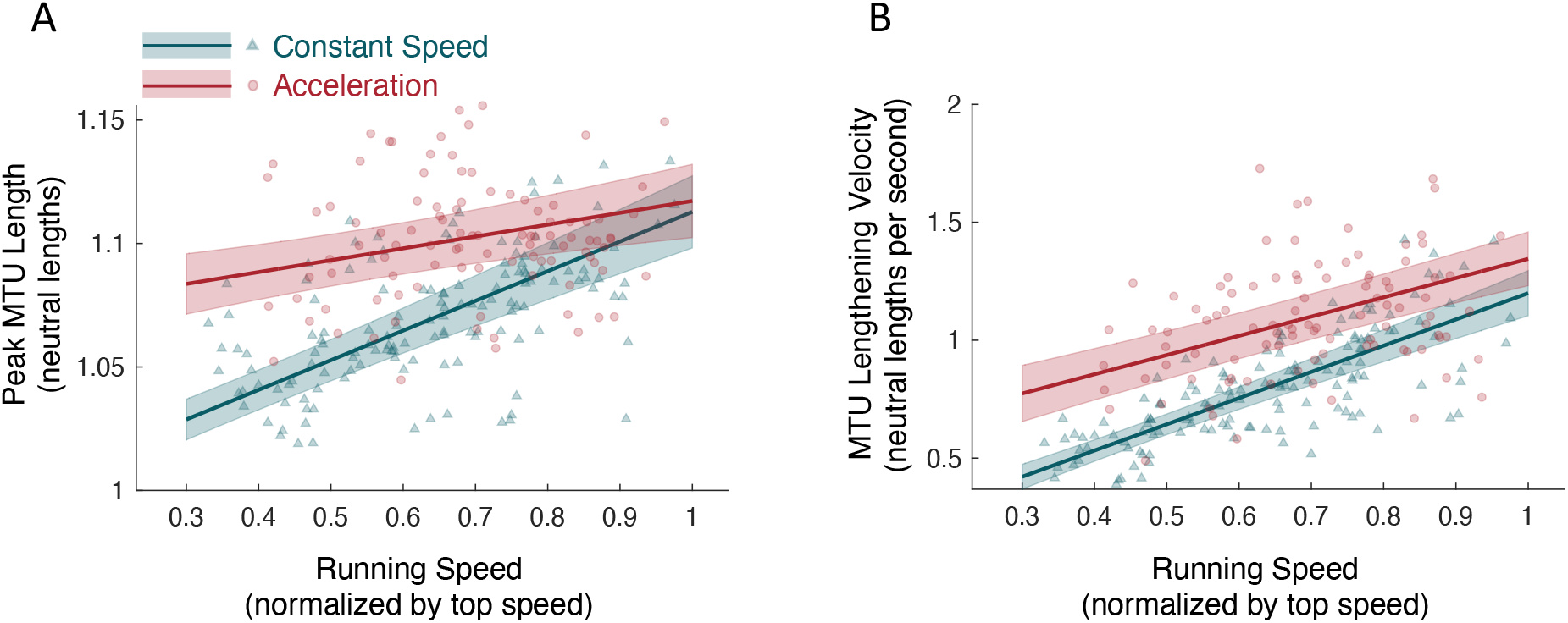
Peak biceps femoris MTU length (A) and average lengthening velocity (B) vs. normalized running speed during both constant speed (teal) and accelerative (red) running. Neutral length is the MTU length in the neutral configuration (upright standing posture). Each data point (triangles for constant speed, circles for acceleration) represents a single step. Solid lines depict the linear mixed model regression lines, and the shaded area illustrates the 90% prediction interval.

The greater peak MTU lengths in accelerative running resulted from a more flexed hip (hip flexion lengthens the hamstrings), despite a more flexed knee (knee flexion shortens the hamstrings) compared to constant speed running (Figure 3). The MTU lengthening effect of hip flexion was greater than the shortening effect of knee flexion because the hamstring moment arm about the hip is greater than the knee. The differences between accelerative and constant speed running in the hip and knee flexion angle at the instant of peak MTU length were greater for slower running speeds (Figure 1, middle and bottom rows). For example, the hip was more flexed (p<0.001) in accelerative than in constant speed running at speeds below 96% of top speed (based on overlap of the 90% prediction intervals, Figure 3A). This greater hip flexion angle at the instant of peak MTU length was due primarily to a more horizontal thigh (Figure 4B) in addition to a more anteriorly tilted pelvis (Figure 4A). Based on the linear mixed model, the hip was flexed 6° more in accelerative running than in constant speed running with every 10% decrease in running speed, where 4° of the difference was due to a more horizontal thigh and 2° to a more anteriorly tilted pelvis.

**Figure 3.**
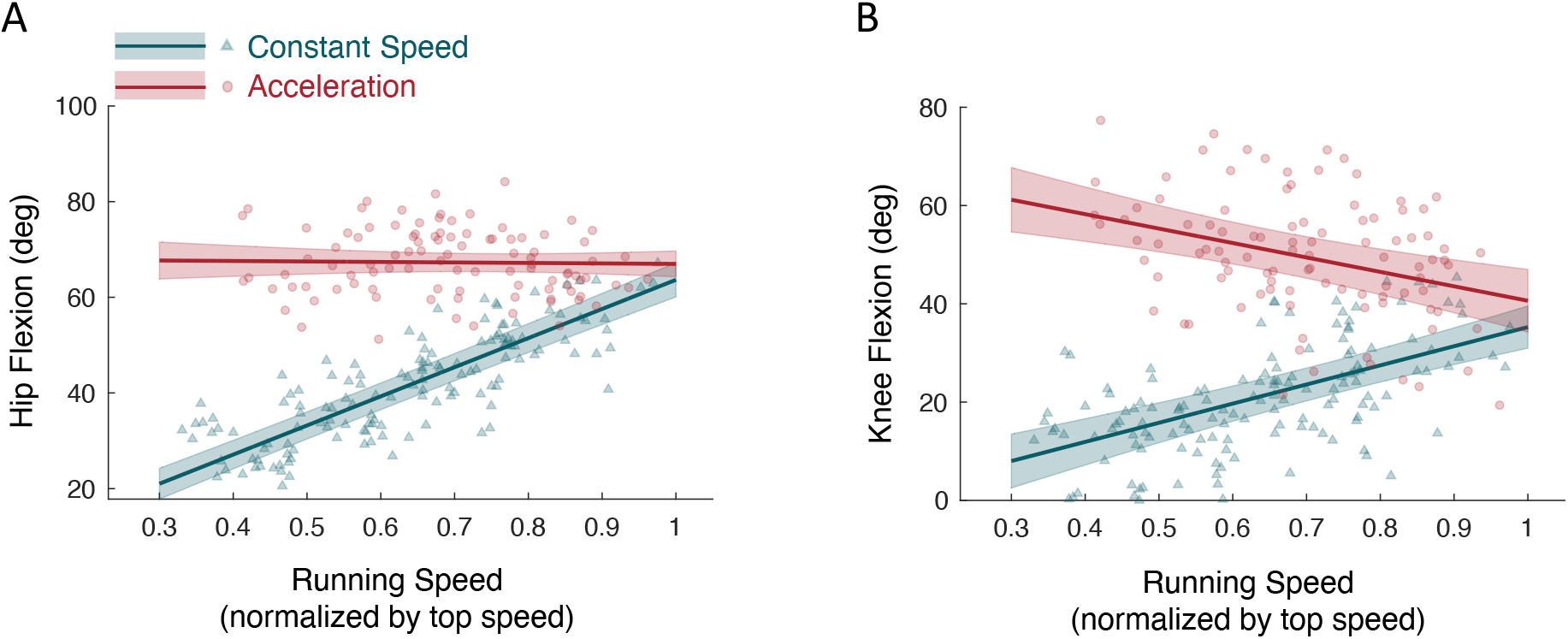
Hip (A) and knee (B) flexion angle at the instant of peak MTU length vs. normalized running speed during both constant speed (teal) and accelerative (red) running. Each data point (triangles for constant speed, circles for acceleration) represents a single step. Solid lines depict the linear mixed model regression lines, and the shaded area illustrates the 90% prediction interval.

**Figure 4.**
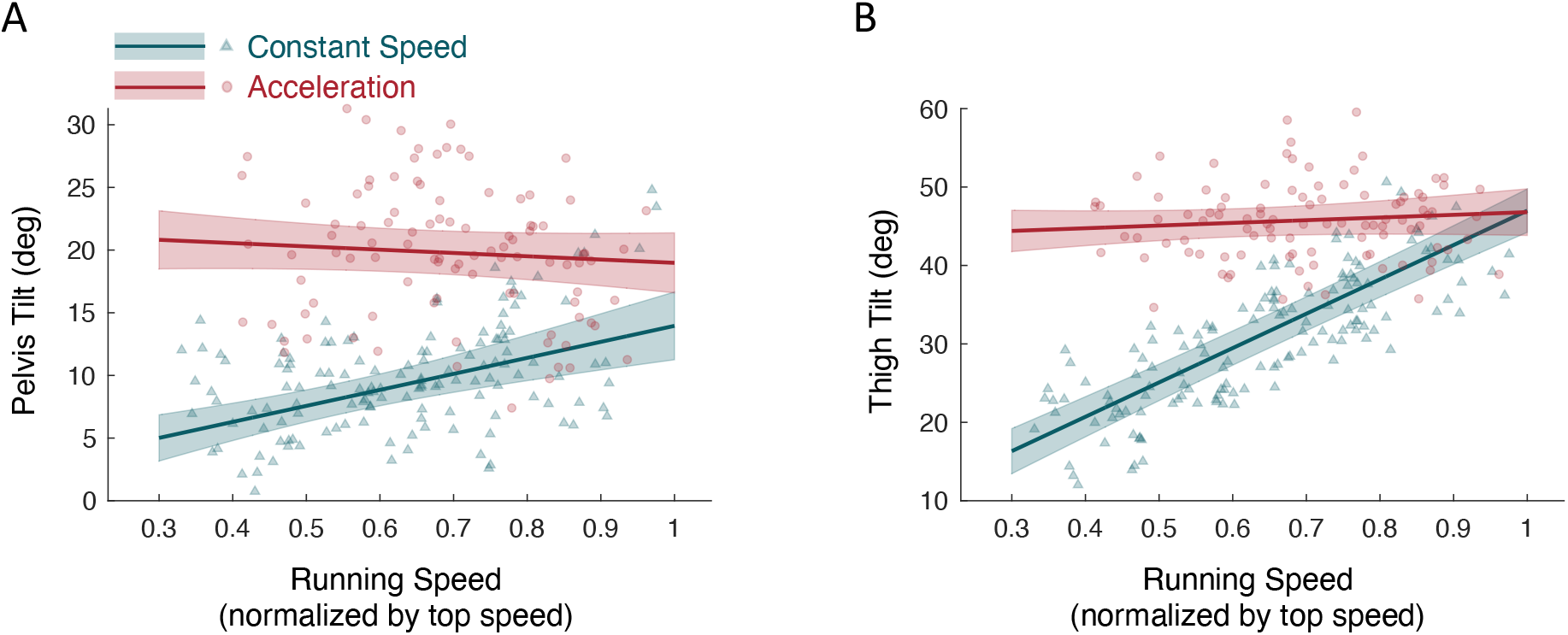
Pelvis (A) and thigh (B) tilt angle at the instant of peak MTU length vs. normalized running speed during both constant speed (teal) and accelerative (red) running. Positive pelvis and thigh tilt angles both contribute to hip flexion. For example, at top speed (normalized running speed = 1) in accelerative running, the pelvis (19°) and thigh (47°) tilt angles sum to 66°, the approximate hip flexion angle in the same condition (Figure 3A). Each data point (triangles for constant speed, circles for acceleration) represents a single step. Solid lines depict the linear mixed model regression lines, and the shaded area illustrates the 90% prediction interval.

The average MTU lengthening velocity was greater during accelerative than in constant speed running across all running speeds (Figure 2B). This resulted from a greater hip flexion speed, rather than from a greater knee extension speed (Figure 5). In fact, below 78% of top running speed, the MTU lengthening velocity resulting from the knee extension speed was greater in constant speed than in accelerative running (based on 90% prediction interval overlap, Figure 5B).

**Figure 5.**
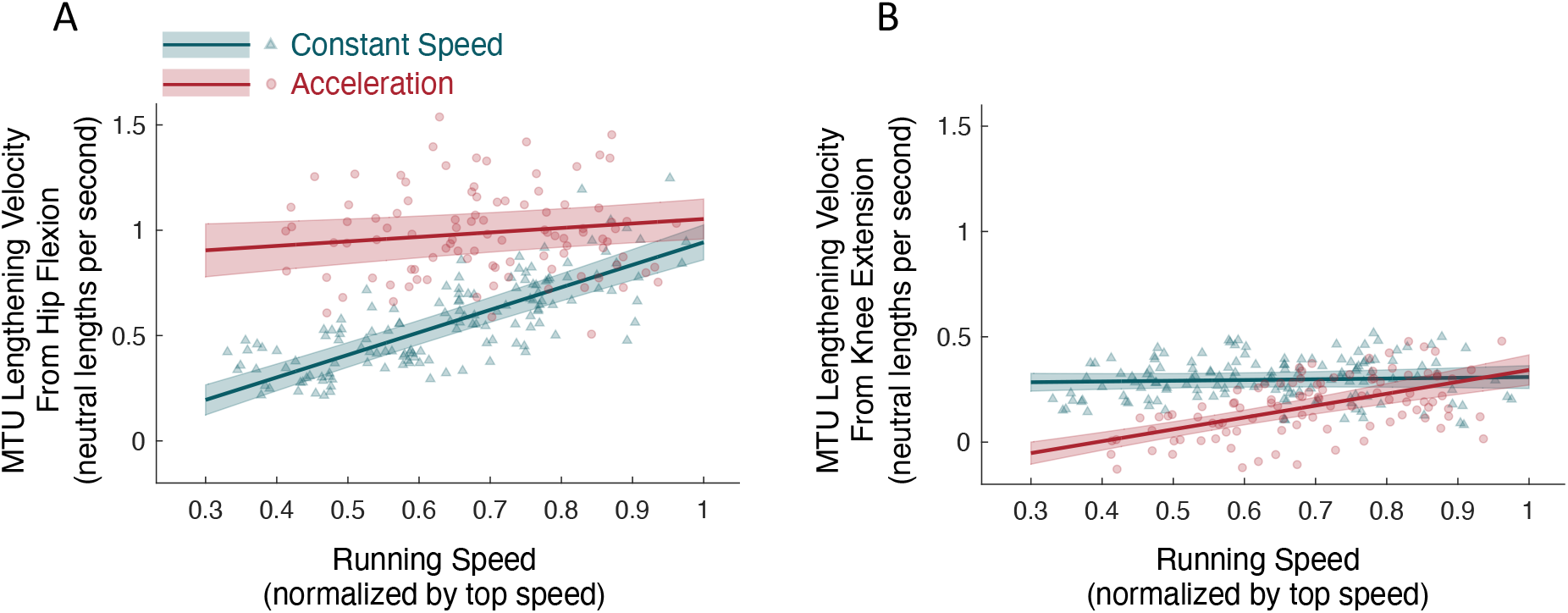
Hip flexion (A) and knee extension (B) speed contributions to the average MTU lengthening velocity vs. normalized running speed during both constant speed (teal) and accelerative (red) running. Neutral length is the MTU length in the neutral configuration (upright standing posture). Each data point (triangles for constant speed, circles for acceleration) represents a single step. Solid lines depict the linear mixed model regression lines, and the shaded area illustrates the 90% prediction interval.

## 4. Discussion

Our results demonstrate that the hamstrings are stretched more and faster during accelerative compared to constant speed running. Furthermore, the differences were more pronounced when accelerating from slower running speeds. This provides a biomechanical explanation for the observation that hamstring injuries often occur when athletes are accelerating^31^ (and not necessarily at high running speeds^22^). Our results also have implications for athlete monitoring. For example, while our data support the use of exposure to high running speeds (e.g., from body-worn GPS units) as a proxy for increased hamstring length and lengthening velocity, they also suggest the accelerative nature of running should be included to more comprehensively characterize exposure to high-risk circumstances.

Our results illustrate how hip-knee coordination underlies the differences in accelerative and constant speed running and the implications for exposing the hamstrings to long lengths. The kinematics of accelerative running were such that the peak MTU length occurred at an average hip flexion angle of 67° (Figure 3A), and this was not influenced by running speed. However, the knee flexion angle at which the peak MTU length occurred was influenced by running speed.

For example, when accelerating from slower running speeds, the knee was more flexed at the instant of peak MTU length than in speed-matched, constant speed running (approximately 3° more knee flexion with every 10% of top speed decrease in running speed). Without this increase in knee flexion, the peak MTU length would have been even greater when accelerating from slower running speeds (i.e., when the capacity to accelerate is greater). Thus, a coordinated motion wherein the hip and knee flex together during the late swing phase of accelerative running reduces the peak length of the hamstrings. In contrast, an accelerative running pattern wherein the knee does not flex synchronously with the hip exposes the hamstrings to longer peak lengths.

The importance of hip-knee coordination in accelerative running is further supported by the observed MTU lengthening velocities. Large hip flexion speeds drove the increase in MTU lengthening velocity during accelerative running compared to constant speed running. This was despite the knee extension speed contributing less to MTU lengthening velocity in accelerative compared to constant speed running (Figure 5B). In fact, our linear mixed models predicted that when accelerating from below approximately 40% of top speed, the knee contributed a net average *shortening* velocity (negative vertical axis in Figure 5B). Therefore, a lack of simultaneous hip and knee flexion during accelerative running would be two-fold detrimental. It would result in greater peak MTU length and greater MTU lengthening velocity. An altered hip-knee coordination in this way is generally consistent with others who have noted the potential influence of altered coordination on injury risk^30^.

Cumulative eccentric MTU work has been proposed as a predictor of muscle damage and would be a candidate proxy for hamstring injury risk. Eccentric work can be interpreted as the average MTU lengthening velocity weighted by MTU force. This supports our use of average MTU lengthening velocity as a kinematic surrogate for eccentric work. Nonetheless, future work should compare muscle kinetics between accelerative running and constant speed running. OpenCap allowed us to quantify the kinematics of accelerative running overground in outdoor conditions. Its use of computer vision presents some limitations, especially regarding measurement of the pelvis tilt angle. The OpenPose pose detection algorithm only identifies 2 key points on the pelvis (the 2 hip joint centers) rendering the pelvis tilt angle unobservable. Observability is made possible following prediction of the positions of additional pelvis-fixed anatomical landmarks from the OpenCap neural network, which is trained on a large dataset that includes running motions. Confidence in our results is supported by similarity with previous studies. Thelen et al. (2005) reported peak MTU lengths between 1.09-1.10 neutral lengths that occurred at hip and knee flexion angles between 63-65° and 44-45°, respectively, during constant speed running at 80-100% of top speed^11^. Under the same conditions, the linear mixed models based on our data predict similar results for peak MTU length (1.09-1.11 neutral lengths) and hip flexion (between 52-64°) but with less knee flexion (28-35°). We also observed the same general pattern of the MTU length trajectories (e.g., compare the top row of our Figure 1 with the top row of Figure 2 from Thelen et al. (2005)). Schache et al. report peak biceps femoris MTU strain of 12% at top speed^26^ which is comparable to our results (11%). The linear models based on our data predict anterior pelvic tilt angles between 14-19° across both accelerative and constant speed running at top speed, which is comparable to the range reported by previous studies (e.g., 10-20°^46^ and 15-30°^38^). We report hip and knee angle trajectories comparable to Higashihara et al.^39^ with slightly lower peak values which may arise because the average top speeds of the sprinters in their study (9.52 m/s) is higher than ours (7.4 m/s).

We considered only accelerative running where running speed was increased (positive acceleration). However, hamstring injuries have also been observed in decelerative running^22,31^. It is unclear how joint and hamstring mechanics differ between decelerative compared to accelerative or constant speed running and should be examined in future work.

## 5. Conclusion

The hamstrings are stretched to greater peak lengths and are subject to greater lengthening velocities in accelerative than in constant speed running. Greater hip flexion and hip flexion speed during accelerative running was the primary driver of the greater hamstring lengths and velocities. Thus, the accelerative nature of running should be considered in addition to running speed for monitoring athlete exposure to high-risk circumstances. These high-risk circumstances can be mitigated in part by coordinating the hip and knee motions during accelerative running.

## Acknowledgments

This work was supported by the Wu Tsai Human Performance Alliance at Stanford University and the Joe and Clara Tsai Foundation, and the National Institutes of Health through Grant P41EB027060. We thank Julie Muccini for her help running experiments.

## Declaration of Generative AI and AI-Assisted Technologies in the Writing Process

None to declare.

## References

1. Ekstrand J, Hagglund M, Walden M. Injury incidence and injury patterns in professional football: the UEFA injury study. Br J Sports Med. 2011;45(7):553–558. doi:10.1136/bjsm.2009.060582

2. Edouard P, Branco P, Alonso JM. Muscle injury is the principal injury type and hamstring muscle injury is the first injury diagnosis during top-level international athletics championships between 2007 and 2015. Br J Sports Med. 2016;50(10):619–630. doi:10.1136/bjsports-2015-095559

3. Lu Y, Pareek A, Lavoie-Gagne OZ, et al. Machine Learning for Predicting Lower Extremity Muscle Strain in National Basketball Association Athletes. Orthop J Sports Med. 2022;10(7):232596712211117. doi:10.1177/23259671221111742

4. Volpi A, Haselman W, Photopoulos C, Banffy M. Regular-Season Injury Rates in the National Football League After an Attenuated Preseason Due to COVID-19. Orthop J Sports Med. 2022;10(11):232596712211337. doi:10.1177/23259671221133776

5. Brooks JHM, Fuller CW, Kemp SPT, Reddin DB. Incidence, Risk, and Prevention of Hamstring Muscle Injuries in Professional Rugby Union. Am J Sports Med. 2006;34(8):1297–1306. doi:10.1177/0363546505286022

6. Maniar N, Carmichael DS, Hickey JT, et al. Incidence and prevalence of hamstring injuries in field-based team sports: a systematic review and meta-analysis of 5952 injuries from over 7 million exposure hours. Br J Sports Med. 2023;57(2):109–116. doi:10.1136/bjsports-2021-104936

7. Ekstrand J, Bengtsson H, Waldén M, Davison M, Khan KM, Hägglund M. Hamstring injury rates have increased during recent seasons and now constitute 24% of all injuries in men’s professional football: the UEFA Elite Club Injury Study from 2001/02 to 2021/22. Br J Sports Med. 2023;57(5):292–298. doi:10.1136/bjsports-2021-105407

8. Martin RL, Cibulka MT, Bolgla LA, et al. Hamstring Strain Injury in Athletes: Clinical Practice Guidelines Linked to the International Classification of Functioning, Disability and Health From the Academy of Orthopaedic Physical Therapy and the American Academy of Sports Physical Therapy of the American Physical Therapy Association. J Orthop Sports Phys Ther. 2022;52(3):CPG1–CPG44. doi:10.2519/jospt.2022.0301

9. Visser H de, Reijman M, Heijboer MP, Bos PK. Risk factors of recurrent hamstring injuries: a systematic review. Br J Sports Med. 2012;46(2):124–130. doi:10.1136/bjsports-2011-090317

10. Fiorentino NM, Rehorn MR, Chumanov ES, Thelen DG, Blemker SS. Computational Models Predict Larger Muscle Tissue Strains at Faster Sprinting Speeds. Med Sci Sports Exerc. 2014;46(4):776–786. doi:10.1249/MSS.0000000000000172

11. Thelen DG, Chumanov ES, Hoerth DM, et al. Hamstring Muscle Kinematics during Treadmill Sprinting: Med Sci Sports Exerc. 2005;37(1):108–114. doi:10.1249/01.MSS.0000150078.79120.C8

12. Chumanov ES, Heiderscheit BC, Thelen DG. The effect of speed and influence of individual muscles on hamstring mechanics during the swing phase of sprinting. J Biomech. 2007;40(16):3555–3562. doi:10.1016/j.jbiomech.2007.05.026

13. Fiorentino NM, Epstein FH, Blemker SS. Activation and aponeurosis morphology affect in vivo muscle tissue strains near the myotendinous junction. J Biomech. 2012;45(4):647–652. doi:10.1016/j.jbiomech.2011.12.015

14. Brooks SV, Zerba E, Faulkner JA. Injury to muscle fibres after single stretches of passive and maximally stimulated muscles in mice. J Physiol. 1995;488(2):459–469. doi:10.1113/jphysiol.1995.sp020980

15. Mair SD, Seaber AV, Glisson RR, Garrett WE. The Role of Fatigue in Susceptibility to Acute Muscle Strain Injury. Am J Sports Med. 1996;24(2):137–143. doi:10.1177/036354659602400203

16. Butterfield TA, Herzog W. Quantification of muscle fiber strain during in vivo repetitive stretch-shortening cycles. J Appl Physiol. 2005;99(2):593–602. doi:10.1152/japplphysiol.01128.2004

17. Danielsson A, Horvath A, Senorski C, et al. The mechanism of hamstring injuries – a systematic review. BMC Musculoskelet Disord. 2020;21(1):641. doi:10.1186/s12891-020-03658-8

18. Yu B, Liu H, Garrett WE. Mechanism of hamstring muscle strain injury in sprinting. J Sport Health Sci. 2017;6(2):130–132. doi:10.1016/j.jshs.2017.02.002

19. Woods C. The Football Association Medical Research Programme: an audit of injuries in professional football--analysis of hamstring injuries. Br J Sports Med. 2004;38(1):36–41. doi:10.1136/bjsm.2002.002352

20. Heiderscheit BC, Hoerth DM, Chumanov ES, Swanson SC, Thelen BJ, Thelen DG. Identifying the time of occurrence of a hamstring strain injury during treadmill running: A case study. Clin Biomech. 2005;20(10):1072–1078. doi:10.1016/j.clinbiomech.2005.07.005

21. Schache AG, Wrigley TV, Baker R, Pandy MG. Biomechanical response to hamstring muscle strain injury. Gait Posture. 2009;29(2):332–338. doi:10.1016/j.gaitpost.2008.10.054

22. Kerin F, Farrell G, Tierney P, McCarthy Persson U, De Vito G, Delahunt E. Its not all about sprinting: mechanisms of acute hamstring strain injuries in professional male rugby union—a systematic visual video analysis. Br J Sports Med. 2022;56(11):608–615. doi:10.1136/bjsports-2021-104171

23. Beato M, Drust B, Iacono AD. Implementing High-speed Running and Sprinting Training in Professional Soccer. Int J Sports Med. 2021;42(04):295–299. doi:10.1055/a-1302-7968

24. Buckthorpe M, Wright S, Bruce-Low S, et al. Recommendations for hamstring injury prevention in elite football: translating research into practice. Br J Sports Med. 2019;53(7):449–456. doi:10.1136/bjsports-2018-099616

25. Kenneally-Dabrowski CJB, Brown NAT, Lai AKM, Perriman D, Spratford W, Serpell BG. Late swing or early stance? A narrative review of hamstring injury mechanisms during high-speed running. Scand J Med Sci Sports. 2019;29(8):1083–1091. doi:10.1111/sms.13437

26. Schache AG, Dorn TW, Blanch PD, Brown NAT, Pandy MG. Mechanics of the Human Hamstring Muscles during Sprinting. Med Sci Sports Exerc. 2012;44(4):647–658. doi:10.1249/MSS.0b013e318236a3d2

27. Miller RH, Umberger BR, Caldwell GE. Limitations to maximum sprinting speed imposed by muscle mechanical properties. J Biomech. 2012;45(6):1092–1097. doi:10.1016/j.jbiomech.2011.04.040

28. Chumanov ES, Heiderscheit BC, Thelen DG. Hamstring Musculotendon Dynamics during Stance and Swing Phases of High-Speed Running. Med Sci Sports Exerc. 2011;43(3):525–532. doi:10.1249/MSS.0b013e3181f23fe8

29. Haralabidis N, Serrancolí G, Colyer S, Bezodis I, Salo A, Cazzola D. Three-dimensional data-tracking simulations of sprinting using a direct collocation optimal control approach. PeerJ. 2021;9:e10975. doi:10.7717/peerj.10975

30. Bramah C, Mendiguchia J, Dos’Santos T, Morin JB. Exploring the Role of Sprint Biomechanics in Hamstring Strain Injuries: A Current Opinion on Existing Concepts and Evidence. Sports Med. Published online September 19, 2023. doi:10.1007/s40279-023-01925-x

31. Aiello F, Di Claudio C, Fanchini M, et al. Do non-contact injuries occur during high-speed running in elite football? Preliminary results from a novel GPS and video-based method. J Sci Med Sport. 2023;26(9):465–470. doi:10.1016/j.jsams.2023.07.007

32. Ekstrand J, Waldén M, Hägglund M. Hamstring injuries have increased by 4% annually in men’s professional football, since 2001: a 13-year longitudinal analysis of the UEFA Elite Club injury study. Br J Sports Med. 2016;50(12):731–737. doi:10.1136/bjsports-2015-095359

33. Barnes C, Archer D, Hogg B, Bush M, Bradley P. The Evolution of Physical and Technical Performance Parameters in the English Premier League. Int J Sports Med. 2014;35(13):1095–1100. doi:10.1055/s-0034-1375695

34. Nagahara R, Matsubayashi T, Matsuo A, Zushi K. Kinematics of transition during human accelerated sprinting. Biol Open. 2014;3(8):689–699. doi:10.1242/bio.20148284

35. Pandy MG, Lai AKM, Schache AG, Lin Y. How muscles maximize performance in accelerated sprinting. Scand J Med Sci Sports. 2021;31(10):1882–1896. doi:10.1111/sms.14021

36. Morin JB, Gimenez P, Edouard P, et al. Sprint Acceleration Mechanics: The Major Role of Hamstrings in Horizontal Force Production. Front Physiol. 2015;6. doi:10.3389/fphys.2015.00404

37. Rabita G, Dorel S, Slawinski J, et al. Sprint mechanics in world-class athletes: a new insight into the limits of human locomotion. Scand J Med Sci Sports. 2015;25(5):583–594. doi:10.1111/sms.12389

38. Schuermans J, Van Tiggelen D, Palmans T, Danneels L, Witvrouw E. Deviating running kinematics and hamstring injury susceptibility in male soccer players: Cause or consequence? Gait Posture. 2017;57:270-277. doi:10.1016/j.gaitpost.2017.06.268

39. Higashihara A, Nagano Y, Ono T, Fukubayashi T. Differences in hamstring activation characteristics between the acceleration and maximum-speed phases of sprinting. J Sports Sci. 2018;36(12):1313–1318. doi:10.1080/02640414.2017.1375548

40. Uhlrich SD, Falisse A, Kidziński Ł, et al. OpenCap: Human movement dynamics from smartphone videos. Marsden AL, ed. PLOS Comput Biol. 2023;19(10):e1011462. doi:10.1371/journal.pcbi.1011462

41. Lai AKM, Arnold AS, Wakeling JM. Why are Antagonist Muscles Co-activated in My Simulation? A Musculoskeletal Model for Analysing Human Locomotor Tasks. Ann Biomed Eng. 2017;45(12):2762–2774. doi:10.1007/s10439-017-1920-7

42. Cao Z, Hidalgo G, Simon T, Wei SE, Sheikh Y. OpenPose: Realtime Multi-Person 2D Pose Estimation Using Part Affinity Fields. IEEE Trans Pattern Anal Mach Intell. 2021;43(1):172–186. doi:10.1109/TPAMI.2019.2929257

43. Ben Mansour K, Rezzoug N, Gorce P. Analysis of several methods and inertial sensors locations to assess gait parameters in able-bodied subjects. Gait Posture. 2015;42(4):409–414. doi:10.1016/j.gaitpost.2015.05.020

44. Seth A, Hicks JL, Uchida TK, et al. OpenSim: Simulating musculoskeletal dynamics and neuromuscular control to study human and animal movement. Schneidman D, ed. PLOS Comput Biol. 2018;14(7):e1006223. doi:10.1371/journal.pcbi.1006223

45. Robertson DGE, Dowling JJ. Design and responses of Butterworth and critically damped digital filters. J Electromyogr Kinesiol. 2003;13(6):569–573. doi:10.1016/S1050-6411(03)00080-4

46. Mendiguchia J, Castaño-Zambudio A, Jimenez-Reyes P, et al. Can we modify maximal speed running posture? Implications for performance and hamstring injuries management. Int J Sports Physiol Perform. 2021;17(3):374–383. doi:10.1123/ijspp.2021-0107

