## Supplementary Material for "Hamstrings are stretched more and faster during accelerative running compared to speed-matched constant speed running"

Table S1. Linear mixed model fixed effects coefficients. Significant ($p<0.05$) differences are bolded. Intercept units for peak MTU length are in neutral lengths, for MTU lengthening velocity and MTU lengthening velocity from knee and hip are in neutral lengths per second, and for hip and knee flexion and pelvis and thigh tilt angle at peak MTU length are in degrees. Slope units are the intercept units per normalized running speed. As noted in the main text, running speeds were shifted for model fitting such that the intercept corresponded to the response at top speed. Thus, for example, the model for peak MTU length during constant speed running is $y = 1.11 + 0.12(x-1)$.

| | Intercept | | | Slope | | |
|---|---|---|---|---|---|---|
| Response | Constant | Accel. | Difference | Constant | Accel. | Difference |
| Peak MTU Length | 1.11 | 1.12 | 0.00 (p=0.431) | 0.12 | 0.05 | **-0.07 (p<0.001)** |
| MTU Lengthening Velocity | 1.20 | 1.35 | **0.15 (p=0.012)** | 1.11 | 0.82 | **-0.30 (p=0.045)** |
| Hip Flex. at Peak Length | 63.65 | 66.98 | 3.33 (p=0.079) | 60.90 | -1.02 | **-61.92 (p<0.001)** |
| Pelvis Tilt at Peak Length | 13.96 | 18.98 | **5.03 (p=0.003)** | 12.77 | -2.63 | **-15.40 (p<0.001)** |
| Thigh Tilt at Peak Length | 46.98 | 46.80 | -0.19 (p=0.906) | 43.80 | 3.42 | **-40.37 (p<0.001)** |
| Knee Flex. at Peak Length | 35.26 | 40.62 | 5.36 (p=0.088) | 38.95 | -29.37 | **-68.32 (p<0.001)** |
| MTU Velocity from Knee | 0.31 | 0.34 | 0.03 (p=0.291) | 0.04 | 0.56 | **0.53 (p<0.001)** |
| MTU Velocity from Hip | 0.94 | 1.05 | **0.11 (p=0.023)** | 1.07 | 0.21 | **-0.86 (p<0.001)** |

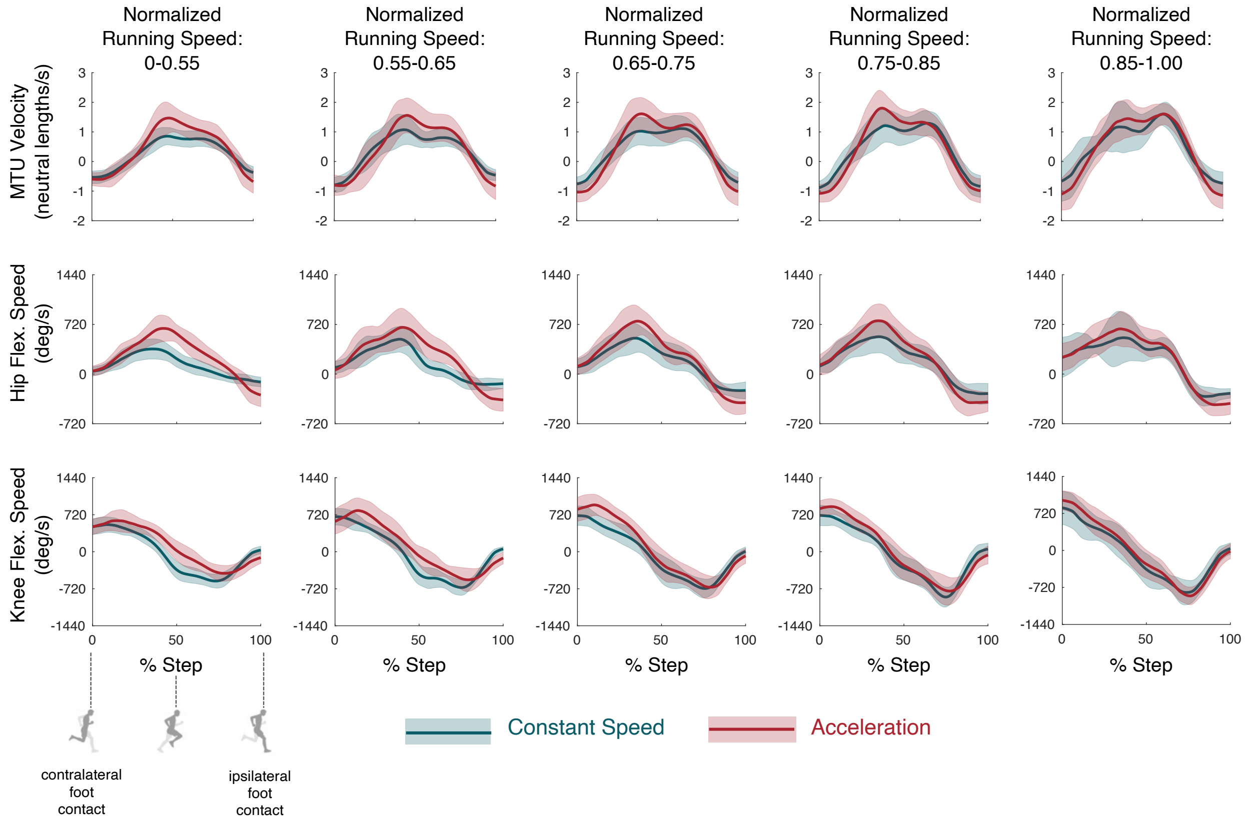


Figure S1. Biceps femoris MTU velocity (top row, units: neutral lengths per second), hip flexion speed (middle row, units: degrees per second), and knee flexion speed (bottom row, units: degrees per second) as a percentage of the step duration for the accelerative (red) and constant speed (teal) running conditions. Solid lines and shaded areas indicate the ensemble mean +/- standard deviation across all participants within the same speed bin (indicated at the top of each column). Running speeds are normalized by the subject-specific top speed. Positive MTU velocities are lengthening velocities (negative are shortening velocities). Positive hip and knee flexion speeds are flexion speeds (negative values are extension speeds).

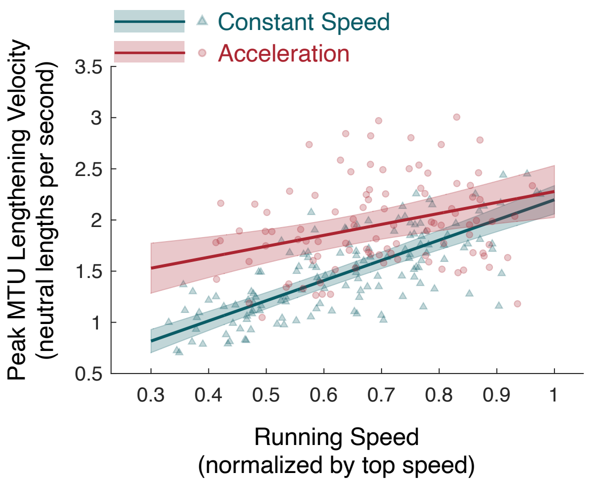


Figure S2. Peak biceps femoris MTU velocity vs. normalized running speed during both constant speed (teal) and accelerative (red) running. Neutral length is the MTU length in neutral configuration (upright standing posture). Each data point (triangles for constant speed, circles for acceleration) represents a single step. Solid lines depict the linear mixed model regression lines, and the shaded area illustrates the 90% prediction interval. The slope and intercept of the linear mixed models for both accelerative and constant speed running are shown in the figure. While the peak lengthening velocity at top speed (intercept) was not significantly different ($p=0.54$) between conditions (2.20 and 2.28 neutral lengths per second for constant speed and accelerative running, respectively), the slope of the accelerative model (1.07 neutral lengths per second per normalized running speed) was significantly less ($p<0.01$) than the constant speed model (1.97 neutral lengths per second per normalized running speed).
